# Evaluation of dsRNA delivery methods for targeting macrophage migration inhibitory factor MIF in RNAi-based aphid control

**DOI:** 10.1101/2021.02.24.432707

**Authors:** Shaoshuai Liu, Maria Jose Ladera-Carmona, Minna M. Poranen, Aart J.E. van Bel, Karl-Heinz Kogel, Jafargholi Imani

## Abstract

Macrophage migration inhibitory factors (MIF) are multifunctional proteins regulating major processes in mammals, including activation of innate immune responses. In invertebrates, MIF proteins participate in the modulation of host immune responses when secreted by parasitic organisms, such as aphids. In this study, we assessed the possibility to use *MIF* genes as targets for RNA interference (RNAi)-based control of the grain aphid *Sitobion avenae* (*Sa*) on barley (*Hordeum vulgare*). When nymphs were fed on artificial diet containing double-stranded (ds)RNAs (*SaMIF*-dsRNAs) that target sequences of the three *MIF* genes *SaMIF1, SaMIF2* and *SaMIF3*, they showed higher mortality rates and these rates correlated with reduced *MIF* transcript levels as compared to the aphids feeding on artificial diet containing a control dsRNA (*GFP*-dsRNA). Comparison of different feeding strategies showed that nymphs’ survival was not altered when they fed from barley seedlings sprayed with *SaMIF*-dsRNAs, suggesting they did not effectively take up dsRNA from the sieve tubes of these plants. Furthermore, aphids’ survival was also not affected when the nymphs fed on leaves supplied with dsRNA via basal cut ends of barley leaves. Consistent with this finding, the use of sieve-tube-specific YFP-labeled Arabidopsis reporter lines confirmed that fluorescent 21 nt dsRNA_Cy3_ supplied via petioles co-localized with xylem structures, but not with phloem tissue. Our results suggest that *MIF* genes are a potential target for insect control and also imply that application of naked dsRNA to plants for aphid control is inefficient. More efforts should be put into the development of effective dsRNA formulations.

## Introduction

Macrophage migration inhibitory factors (MIFs) are multifunctional proteins regulating major processes in mammals, including activation of innate immune responses (Mitchell and Bucala 2000). MIF proteins also play a role in innate immunity of invertebrates and participate in the modulation of host immune responses when secreted by parasitic organisms such as aphids (Rosani et al. 2019; Ghosh et al. 2020). A broad survey of the presence of *MIF* genes across 803 species of plants, fungi, protists, and animals identified them in all eukaryotes. MIFs seem to be essential and highly conserved in some kingdoms (e.g., plants), while they appear more dispensable in other kingdoms (e.g., in fungi) or present in several diverged variants (e.g., insects), suggesting potential neofunctionalizations within the protein superfamily (Michelet et al. 2019). MIFs were discovered in 1966 as a product of activated T cells that limited the random migration of macrophages *in vitro* (David 1966). Subsequently, it was shown that MIFs not only are involved in cell proliferation and apoptosis but play a vital role in the host response against parasitic infection (Calandra and Roger 2003) and vice versa in parasite virulence (Ghosh et al. 2020).

MIFs of aphids also are involved in the response to pathogens and mutualistic symbionts (Dubreuil et al. 2014). Multiple copies of *MIF* genes were found in aphid genomes, including pea aphid (*Acyrthosiphon pisum, Ap)* and green peach aphid (*Myzus persicae, Mp*) (Dubreuil et al., 2014). MIFs are secreted in aphid saliva during feeding, thereby inhibiting major plant immune responses and therefore are crucial to plant infestation (Naessens et al. 2015). Ectopic expression of *MIFs* in leaf tissues inhibited major plant immune responses, such as the expression of defense-related genes, callose deposition, and hypersensitive cell death. Functional complementation analyses showed that MIF1 is the key member of the MIF protein family that allows aphids to exploit their host plants.

Aphids are one of the largest groups of phloem-feeding pests, which can cause huge losses in agriculture and horticulture worldwide (Jaouannet et al. 2014; Pons et al. 2020). They colonize the leaves and stalks, and migrate later towards the ears and settle among the bracts and kernels in the milky-ripe stage of corn plants. A massive withdrawal of sieve-tube components weakens the plant and eventually leads to a reduced overall yield. In most cases, aphids act as important vectors of viruses to spread plant disease (Ng and Perry 2004; Will et al. 2007). More than 5,000 aphid species have been described (The International Aphid Genomics 2010).

We investigated the possibility that *MIF* genes can be used as targets for RNAi-based insect control in plants. Several studies have shown that aphids are sensitive to double-stranded (ds)RNA (Jaubert-Possamai et al. 2007; Pitino et al. 2011) and therefore are amenable to RNAi strategies in crop protection (Christiaens et al. 2020; Liu et al. 2020). In 2015, Abdellatef and colleagues showed that dsRNA derived from the gene encoding Salivary Sheath Protein (SHP), when expressed in barley, strongly reduced the survival of the grain aphid *Sitobion avenae* (*Sa*) (Abdellatef et al. 2015). Similar results were obtained when the green peach aphid was grown on transgenic *Arabidopsis thaliana* expressing dsRNA with homology to the *MpC002* gene (Coleman et al. 2015). The *C002* gene was first described by Mutti et al. (2008) and is predominantly expressed in the salivary glands of aphids.

The degree and the persistence of RNAi in aphids is strong as evidenced by the finding that target genes were also down-regulated in nymphs born from mothers exposed to dsRNA-producing transgenic plants. Notably, *S. avenae* and *M. persicae* aphids reared on transgenic barley (Abdellatef et al. 2015) or Arabidopsis (Coleman et al. 2015), expressing dsRNA against salivary protein components, even showed a decline in survival over several generations. These reports strongly support earlier proposals to use RNAi-based strategies for insect control (Price and Gatehouse 2008; Burand and Hunter 2013).

While transgenic strategies using dsRNA-expressing plants have proven successful in insect control, other strategies might also be applicable. Injection and ingestion of dsRNAs also can induce significant levels of gene silencing in insects (Tomoyasu and Denell 2004; Zhu et al. 2011). Thus, it also might be feasible to deliver dsRNA through foliar application (San Miguel and Scott 2016; Gogoi et al. 2017). The purpose of our study was to assess the potential of *MIF* genes as a target for pest control by oral delivery of dsRNAs derived from gene sequences of three *Sitobion avenae MIF* genes. We also compared the efficiency of different dsRNA delivery strategies, including exposure of aphids to artificial diet vs. leaf spray application and a sucrose-aid delivery in order to provide theoretical support for future application.

## Results

### Prediction of *MIF* genes in *Sitobion avenae* (*Sa*)

Genomic *MIF* sequences of evolutionarily distant species from hemipterans revealed a highly conserved structure (Dubreuil et al. 2014; Michelet et al. 2019). With the aim to deduce *MIF* gene sequences in *Sa* from currently available expressed sequence tags (ESTs) in public databases (https://www.ncbi.nlm.nih.gov/), we searched for *MIF* genes in insect genomes. Based on known peach aphid *Myzus persicae* and pea aphid *Acyrthosiphon pisum* sequence data, partial sequences of *SaMIF1, SaMIF3*, and *SaMIF4* were predicted, amplified by PCR using degenerate primers (Table S1) and sequenced. Sequence alignment, which also included the already published *SaMIF2* sequence (Dubreuil et al. 2014), confirmed that *SaMIFs* are highly conserved in aphids’ evolutionary history (Fig. S1). The identified *SaMIF* sequences (Table S2) were cloned and used as a template for dsRNA production.

### Detection of fluorescence labeled dsRNA in aphids’ midguts after feeding

We conducted dsRNA feeding experiments to assess the effect of *MIF* gene silencing on aphid survival. Since the sucrose concentration in artificial diet is critical, we first tested the optimal concentration of sugar supply. We found that a concentration of 7.5% (w/v) sucrose is optimal for the survival of *Sa* (Fig. S2). Next, we investigated the uptake of fluorescent-labeled dsRNA by *Sa* nymphs from artificial diet. To this end, *SaMIF1*-dsRNA (223 nt; Table S2) labeled with UTP-PEG_5_-AF488 during the dsRNA synthesis was added to the artificial diet at a concentration of 250 ng/μL. A fluorescent signal was observed in the midgut of *Sa* nymphs within 24 h and spread further into the body within 48 h (Fig. 1).

**Fig. 1.**
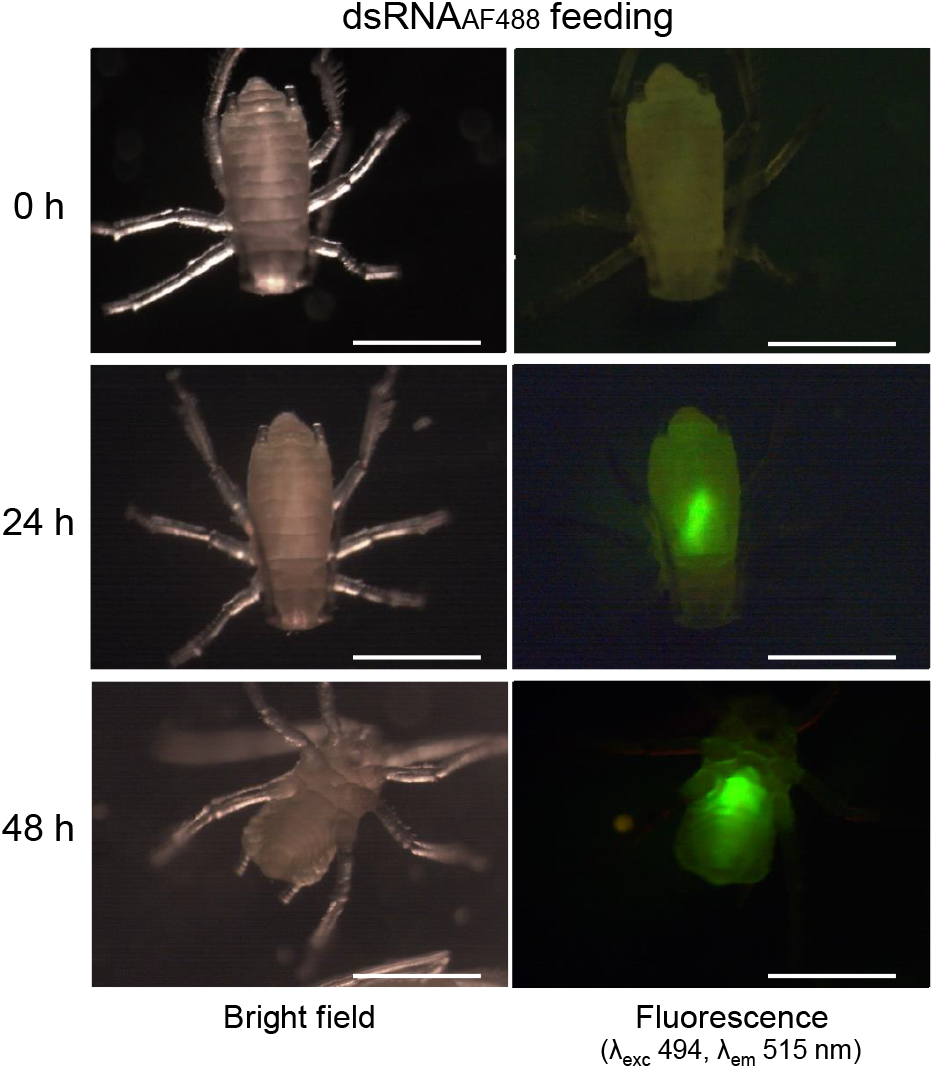
Uptake of fluorescence-labeled dsRNA from an artificial diet and its spreading inside *Sitobion avenae*. Pictures were taken at 0 h, 24 h and 48 h after onset of feeding. The artificial diet contained 250 ng/μL *SaMIF1*-dsRNA_A488_. Fluorescence was detected in the insect gut. Left panels: stereo-microscopic analysis under bright field; right panels: fluorescence stereo microscopic analysis: excitation/emission wavelength (483 nm/506 nm), scale bars = 500 µm.

### The impact of different *SaMIF*-dsRNAs on aphids’ survival

Aphid MIFs are involved in the regulation of plant immune responses, but it remains largely unknown how the respective members of the MIF family contribute to this activity. In *Mp*, mainly MIF1 functions as secreted salivary protein to suppress host immunity (Naessens et al. 2015). We investigated the effect of silencing different *SaMIF* genes on *Sa*’s survival. Since expression of *MIF1, MIF2* and *MIF3* are strongly induced after immune challenge in *Mp* (Dubreuil et al. 2014), we placed our focus on these genes. One-day-old *Sa* nymphs were fed with artificial diet containing dsRNAs directed against *SaMIF1* (*SaMIF1*-dsRNA, 223 nt), *SaMIF2* (*SaMIF2-*dsRNA, 323 nt), *SaMIF3* (*SaMIF3*-dsRNA, 212 nt), and *Green fluorescent protein* (*GFP*-dsRNA, 476 nt) (see Table S1) at two different doses, 250 ng/µL and 125 ng/µL. We found that survival rates of aphids treated with either *SaMIF-*dsRNA *vs. GFP*-dsRNA were significant reduced (Kaplan-Meier analysis and log-rank test, p ≤ 0.0001) at day 4 of feeding with 250 ng/µL (Fig. 2a). Feeding with the lower concentration of *SaMIF*-dsRNAs (125 ng/µL) only resulted in a statistically significant lower survival rate after treatment with *SaMIF1*-dsRNA (Fig. 2b). This finding also confirms that beyond the anticipated function of MIFs as effector interacting with the plant’s defence, MIFs have essential endogenous function in the aphid (Dubreuil et al. 2014).

**Fig. 2.**
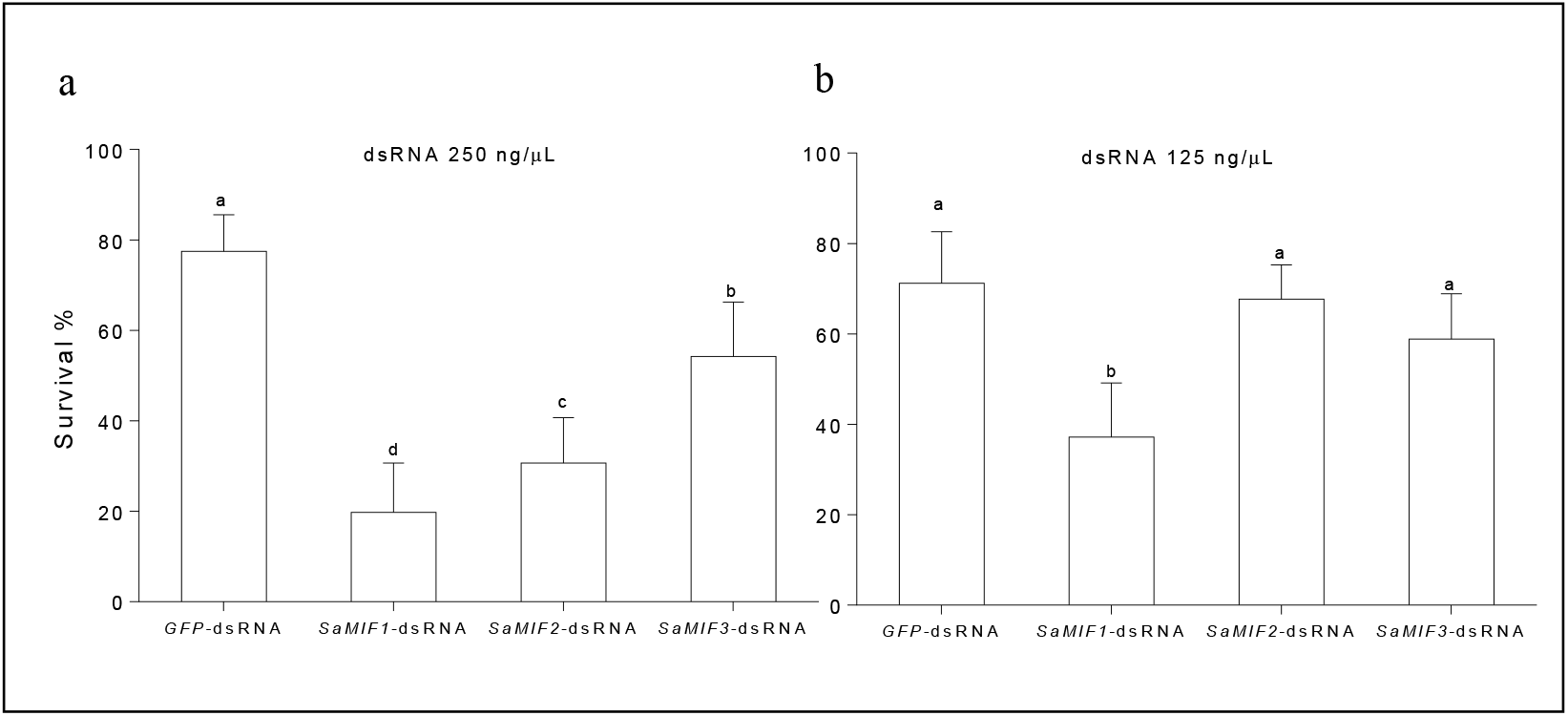
*Sitobion avenae* survival rates after four days of feeding on artificial diet supplied with dsRNA present as percent of control (no dsRNA in the diet). *SaMIF1*-dsRNA, *SaMIF2*-dsRNA, and *SaMIF3*-dsRNA were used with concentration of 250 ng/µL (a) and 125 ng/µL (b). *GFP*-dsRNA was used as an additional control, since a target for this dsRNA is lacking in aphids. Survival data were evaluated by Kaplan-Meier analysis and log-rank test based on three biological replicates. Bars represent means (± SD) from three independent replicates. Different letters indicate significant differences at p < 0.001.

### The impact of *SaMIF*-dsRNA on *MIF* target down regulation

Next, we determined target gene silencing upon feeding aphids with the respective *SaMIF*-dsRNA (250 ng/µL) in artificial diet by 72 h of feeding using RT-qPCR. Consistent with the effects of dsRNA on aphids’ survival, transcript levels of all three *SaMIF* genes were reduced significantly (Student’s *t*-test, p < 0.05) (Fig. 3). These data further substantiate that the effect of *SaMIF*-dsRNAs on *Sa* is based on RNAi-mediated gene silencing.

**Fig. 3.**
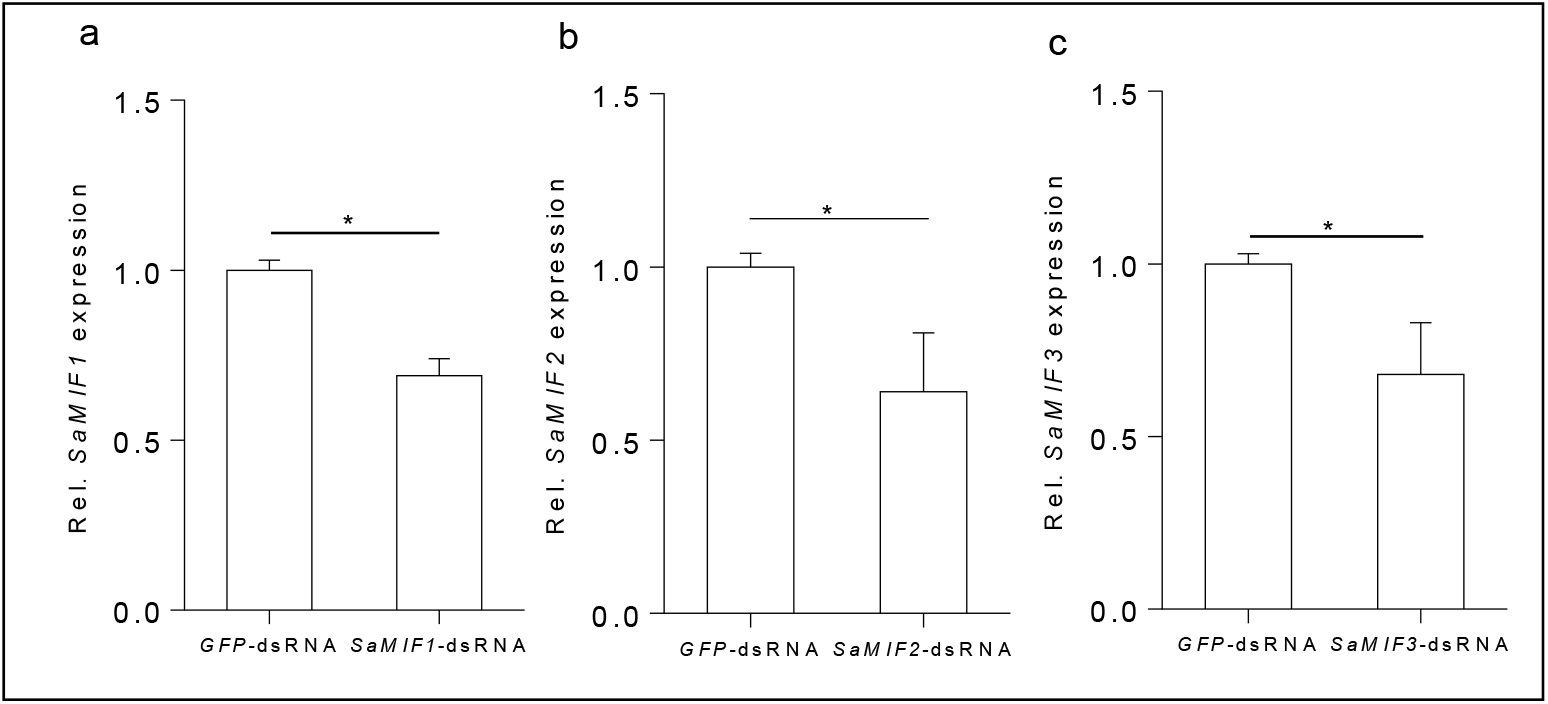
Relative expression of target genes *SaMIF1, SaMIF2* and *SaMIF3* in *Sitobion avenae* fed on an artificial diet containing 250 ng/µL of the respective *SaMIF-*dsRNA. RT-qPCR analysis data for (a) *SaMIF1*, (b) *SaMIF2* and (c) *SaMIF3* were normalized to the aphid’s *Ribosomal protein L27* (*Rpl*27) gene. *GFP-*dsRNA was used as a control. Bars represent means (± SD) from two independent replicates. The asterisks indicate significant differences (Student’s *t*-test; p < 0.05).

### The impact of *SaMIF*-dsRNA mixtures on aphids’ survival

The above data indicate that *SaMIF1* plays a prominent role in the survival of aphids. To further assess *SaMIF1* as a target, we comparatively analysed the effects of *SaMIF1* silencing versus a triple gene silencing of all three *MIF* genes on the survival of *Sa*. Nymphs were treated with *i. SaMIF1*-dsRNA (187.5 ng/µL), *ii*. a mixture of *SaMIF1*-dsRNA, *SaMIF2*-dsRNA, and *SaMIF3*-dsRNA (each at a concentration of 62.5 ng/µL in the artificial diet) and *iii. GFP*-dsRNA (187.5 ng/µL) as control. The relatively low concentration of individual *SaMIF*-dsRNAs in the mixture was chosen because we did no<u>t</u>expect a measurable effect on aphid survival when administered as single dsRNA doses (see Fig. 2). We found that *Sa*’s survival rates treated with either *SaMIF1*-dsRNA or the mixture of *SaMIF-*dsRNAs were significantly reduced (Kaplan-Meier analysis and log-rank test, p ≤ 0.0001) after 5 days as compared with *GFP*-dsRNA treatments (Fig. 4). This suggests that the activity of single dsRNAs are not additive but might have a synergistic effect on the aphid mortality instead.

**Fig. 4.**
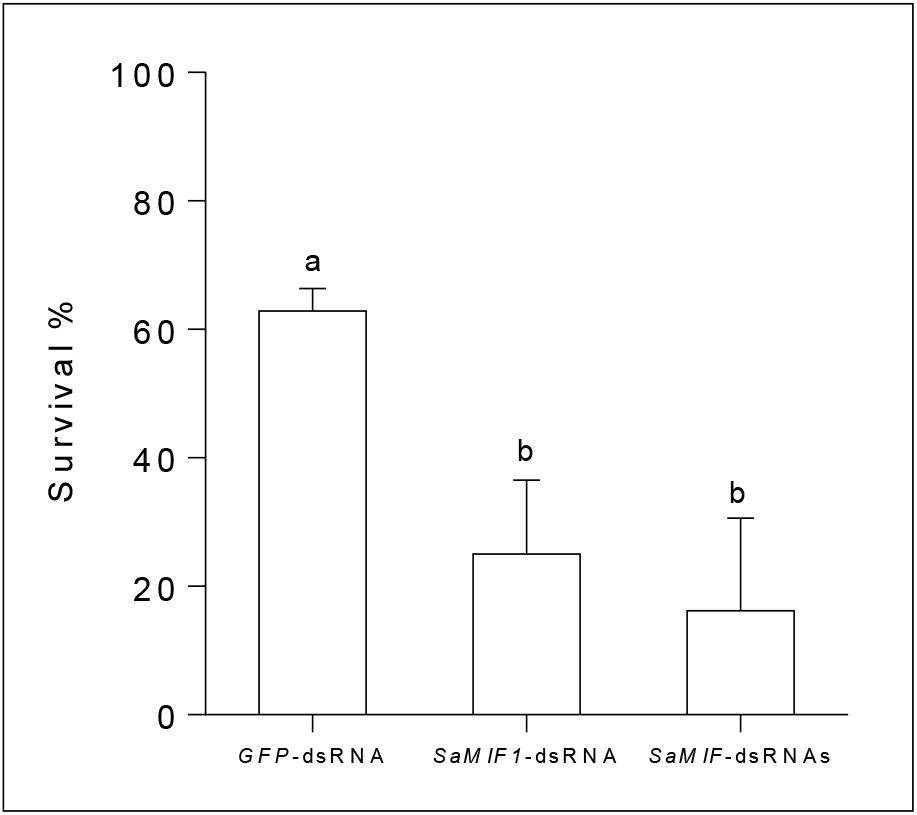
Aphid survival after five days of feeding on artificial diet supplied with *SaMIF1*-dsRNA (187.5 ng/µL), a mixture of *SaMIF1*-dsRNA, *SaMIF2*-dsRNA and *SaMIF3*-dsRNA (each 62.5 ng/µL) or *GFP*-dsRNA (187.5 ng/µL) as a control. Bars represent mean values (± SD) of three biological replicates. Survival data were evaluated by Kaplan-Meier analysis and log-rank test. Different letters indicate significant differences at p < 0.0001.

### *SaMIF1*-dsRNA spray application to barley seedlings had no effect on aphids’ survival

It has been controversially discussed as to whether application of exogenous dsRNA to plants results in its accumulation in the phloem tissue, which is a prerequisite for the RNAi-based control of phloem-feeding insects (Gogoi et al. 2017; Dalakouras et al. 2018). We investigated the possibility that direct application of *SaMIF1*-dsRNA to plants have an effect on *Sa*’s survival, when feding from these plants. Therefore, three barley seedlings per pot were sprayed with 10 µg *SaMIF1*-dsRNA (500 µL of a 20 ng/μL solution), and seedlings, which were infested 24 h later with 50 one-day-old *Sa* nymphs, kept in confined jars (Fig. S3a). Compared to *GFP*-dsRNA-treated control plants, we found no significant differences in the survival rates of aphids feeding on *SaMIF1*-dsRNA-treated plants (Fig. S3b) and controls. This finding implies that spray application to leaves does not result in the accumulation of sufficient amounts of dsRNA or small RNA duplexes derived from it in the sieve tubes and suggests that spray-treatment of naked, unformulated dsRNA probably does not meet the requirements of efficient crop protection.

### Sucrose-aided dsRNA delivery to barley leaves

Next, alternative experimental designs were evaluated for simple and rapid screening of potential dsRNA targets for aphid control. Oligodeoxynucleotide (ODN)-directed gene silencing in barley is mediated by passive vascular feeding of ODN through cut leaves in sucrose solution via co-import of sucrose and negatively charged ODN molecules (Sun et al. 2005), suggesting that ODNs reached the leaf symplast and entered living cells. This report, together with accumulating evidence for xylem-to-phloem solute transport (van Bel 1990) and the presence of exo/endocytosis mechanisms in xylem vessels (Botha et al. 2008; Słupianek et al. 2019), prompted us to investigate whether the cut leaf delivery method could also be used to deliver dsRNA molecules to plant cells, including the phloem tissue. Detached leaves from two-week-old barley seedlings were dipped with the basal end into 1 mL of a solution of 200 mM sucrose and 20 µg *SaMIF1*-dsRNA (Fig. S4a), and kept in the dark for 24 h. As shown in Fig. 5a-c, dsRNA was taken up through the cut ends as revealed by the detection of fluorescence in upper segments of the detached leaves. In barley leaf cross-sections, fluorescence was associated with the vascular bundle, especially the xylem parenchyma cells (Fig. 5d-g). Note that bigger xylem vessels lose their content during preparation of cross-sections due to flushing with the fluid set free by the cut cells and thus do not show fluorescence.

**Fig. 5.**
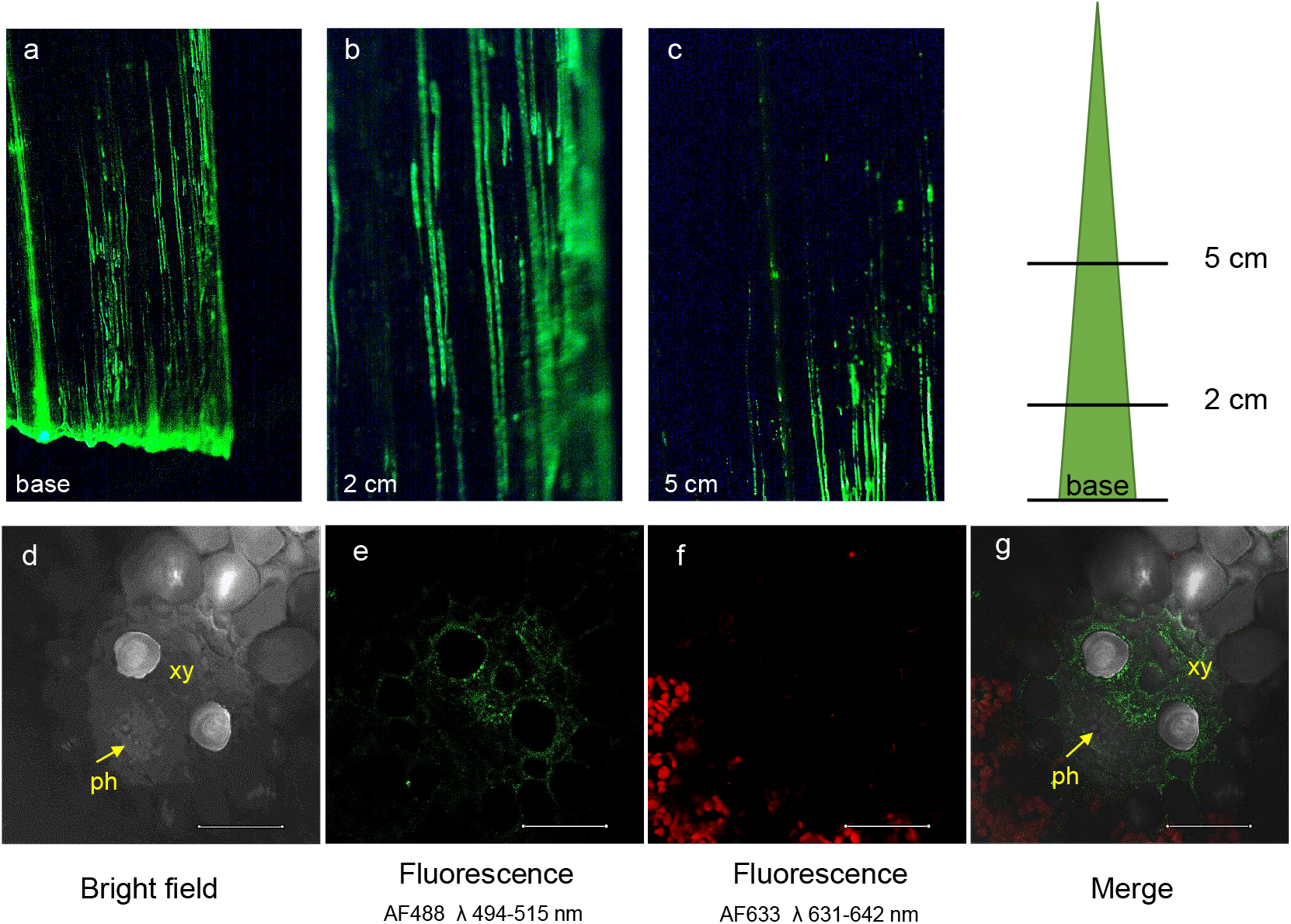
Confocal images of detached barley leaves having absorbed fluorescence-labeled *SaMIF1*-dsRNA_A488_ through cut basal ends. The leaf base was submerged in 1 mL of 200 mM sucrose solution containing 20 µg dsRNA. Surface views of **a**, leaf base; **b**, leaf segment 2 cm away from the base at 24 h after onset of soaking. **c**, leaf segment 5 cm from the base 48 h after onset of soaking. **d-g**, leaf cross-section (3 cm from the cutting), photographs taken at three days after onset of the *SaMIF1*-dsRNA_A488_ treatment. The green color represents the fluorescence (λ_exc_ 494, λ_em_ 515 nm) of the Alexa Flour 488 (AF488) dye. xy, xylem; ph, phloem; bs, bundle sheath.

Next, the survival of *Sa* on *SaMIF1*-dsRNA *vs. GFP*-dsRNA-treated detached barley leaves was recorded after seven days of infestation. Overall, there was no significant difference in the *Sa*’s survival rates on treated and control leaves (Fig. S4b). Consistent with this finding, no difference was found in the expression of the *SaMIF1* target gene in *Sa* fed on *SaMIF1*-dsRNA *vs. GFP*-dsRNA treated leaves (Fig. S4c).

To further substantiate this finding, we conducted the sucrose-aided RNA uptake experiment with *SaSHP*-dsRNA (470 bp; see Table S2), which is known to target the *SaSHP* gene thereby strongly reducing the survival of the aphids on barley (Abdellatef et al. 2015). As shown in Fig. S4b, feeding on *SaSHP*-dsRNA treated leaves also had no effect on aphids’ survival and expression of the *SHP* gene in *Sa* was not affected (Fig. S4d).

### Petiole-mediated uptake of 21 nt dsRNA_Cy3_ in Arabidopsis follows the xylem-route

To further confirm the absence of microscopically detectable exchange of dsRNA between xylem and phloem vessels, when dsRNA is supplied via petioles, we used the Arabidopsis reporter line *Arabidopsis thaliana SUC2::4xYFP*, in which the promoter of the phloem-specific *SUC2* is fused with *Yellow fluorescent protein* (YFP), allowing visualization of the sieve-tubes (Marquès-Bueno et al. 2016). Leaves from thirty-two-day-old plants were inserted with the petioles in nuclease-free water containing fluorescent 21 nt dsRNA_Cy3_ (20 µM). After 24 h, confocal images were taken from different segments of the petioles. We found that dsRNA_Cy3_ was localized in the xylem, and its signal did not overlap with the YFP fluorescence of the phloem (Fig. 6a-h). Moreover, sucrose-aid uptake by petioles resulted in the same localization of Cy3 fluorescence in the xylem vessels (Fig. S5). This result is consistent with our finding that the survival of aphids is not negatively affected when they feed on leaves treated with dsRNA supplied via cut leaf ends. Thus, in contrast to reports showing that ODN can be introduced into plant cells via cut leaf ingestion, our data show that this method of introduction does not result in sufficient uptake of dsRNA or small RNA derivatives to affect aphids or be detected by fluorescence techniques.

**Fig. 6.**
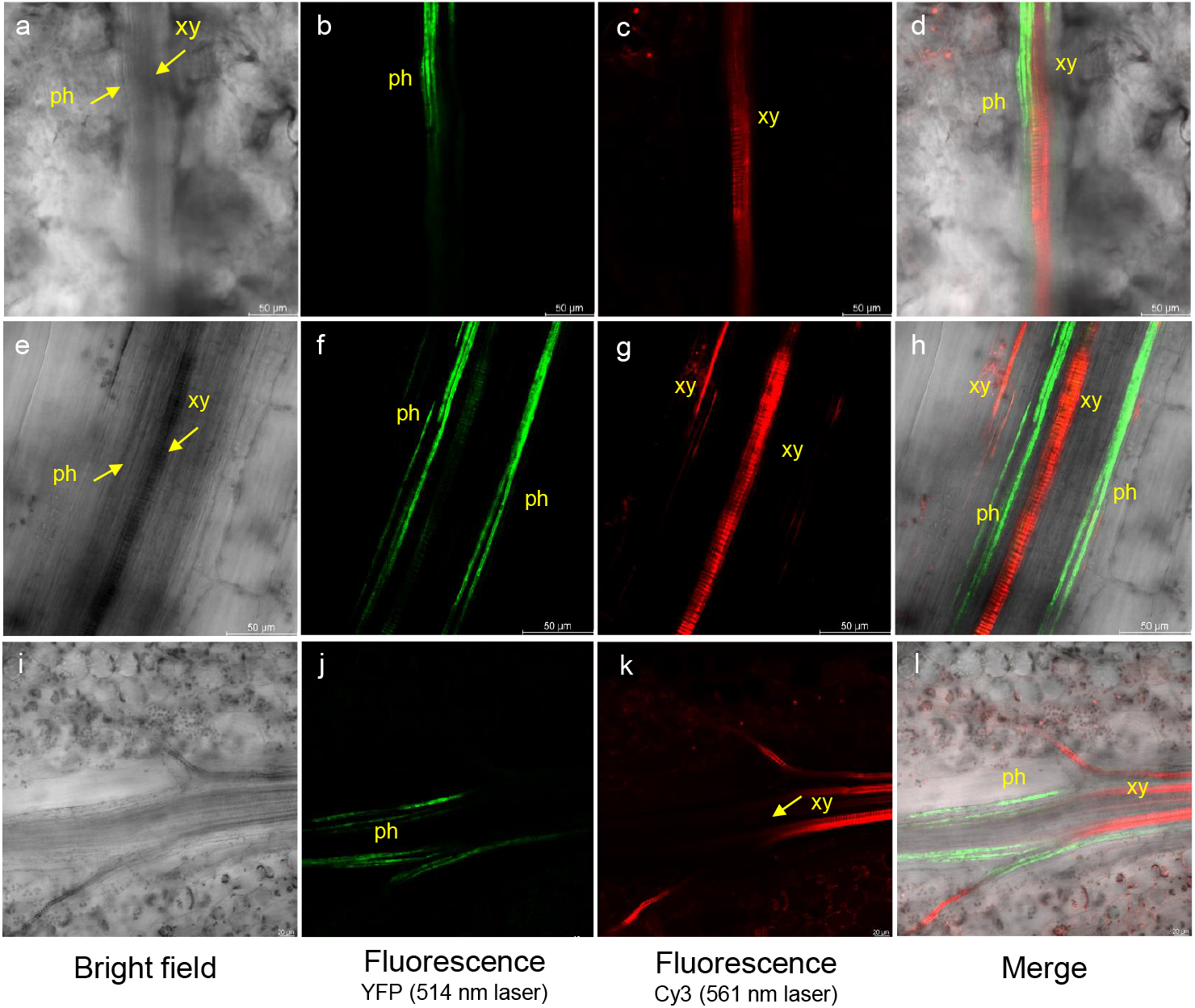
Uptake of labeled dsRNA into *Arabidopsis thaliana* petioles and leaves. Confocal images of the reporter line *SUC2::YFP*. **a-h**, cut petiole ends were submerged for 24 h in nuclease-free water containing 20 µM 21-nt siRNA_Cy3_ and cross-sections were examined at the base (a-d) and in the middle of the petiole (e-h). **i-l**, leaves were dropped with 21 nt dsRNA_Cy3_ (20 µM) for 24 h. Images were taken with a confocal microscope from different segments of the petiole. The red color, which is restricted to the xylem vessels, represents Cy3 fluorescence (λ_exc_555nm, λ_em_569 nm) and the green color represents the phloem-based YFP fluorescence (λ_exc_514 nm, λ_em_527 nm). xy, xylem; ph, phloem.

### dsRNA delivery to leaves also follows the xylem route

Finally, we used the *Arabidopsis thaliana SUC2::4xYFP* reporter line to visualize the uptake of fluorescence dsRNA from the leaf surface (Fig. 6i-l). Arabidopsis leaves were treated with 1 µL drop and four drops per plant of a 20 µM solution of dsRNA_Cy3_. After five days, confocal images were taken from different segments of the leaves. We found that dsRNA_Cy3_ was localized in the xylem, and its signal did not overlap with the YFP fluorescence of the phloem. These finding supports our notion that leaf-applied naked dsRNA does not reach the plant symplast and is therefore an inappropriate method for aphid control.

## Materials and Methods

### Plant Material and Aphids rearing

Spring barley (*Hordeum vulgare* L.) cv. Golden Promise (GP) was used in all experiments. *Arabidopsis thaliana* (Col-0) *SUC2::4xYFP* lines were purchased from NASC (N2106107). Plants were grown under controlled conditions in a climate chamber at 22°C/18°C day/night with 65% relative humidity, a 16 h photoperiod and a photon flux density of 240 μmol m^−2^ s^−1^. Arabidopsis seedlings were grown in vertical plates containing half strength MS medium (Murashige and Skoog 1962), 0.5% of sucrose and 0.7% of agar. The grain aphid (*Sitobion avenae, Sa*) monoclonal population used in this study was reared on three-week-old GP plants in a climate chamber at 22°C with a 16 h photoperiod and a photon flux density of 240 μmol m^−2^ s^−1^. One-day-old fresh synchronized nymphs were used for the experiments (Abdellatef et al. 2015).

### RT-qPCR, Transcript Analysis

RT-qPCR was performed with the Applied Biosystems QuantStudio 5 Real-Time PCR system. Amplifications were performed with SYBR^®^ green JumpStart Taq ReadyMix (Sigma-Aldrich). To quantify the target genes expression, the transcript was normalized with *Ribosomal gene L27* (*RPL27*, NM_001126221.2) (Table S1) (Zhang et al. 2013). The program was performed with 95°C for 5 min, 40 cycles (95°C for 30 sec, 57°C for 30 sec, 72°C for 30 sec. Transcript levels of genes were determined via the 2^−ΔΔ Ct^ method (Livak and Schmittgen 2001) by normalizing to the amount of reference gene transcript.

### dsRNA synthesis

The Si-Fi software was used to select the donor sequences for the RNAi design (Luck et al. 2019). *SaMIF* genes were cloned into pGEM-T-easy vector, using the degenerate primers listed in Table S1, and the resulting plasmids were used as templates for the synthesis of dsRNA. Plasmids pGEM-T-easy-*SHP* and pGEM-T-easy-GFP contain respective *SaSHP* and *GFP* gene sequences (Table S2). The target sequences were amplified from the plasmid DNAs using primers containing T7 polymerase promotor or phi6 polymerase promoter sequences at their 5’-end (Table S1). *SaMIF2-, SaSHP-* and *GFP*-dsRNAs were produced using a single-tube transcription and replication reaction catalyzed by the T7 DNA-dependent RNA polymerase and the phi6 RNA-dependent RNA polymerases (Aalto et al., 2007; Levanova and Poranen, 2018). The produced dsRNAs were enriched using stepwise fractionation with LiCl, followed by precipitation with sodium acetate and thorough washing of the resulting pellet with 70% ethanol. Alternatively, *SaMIF*- and *GFP*-dsRNAs were generated using MEGAscript T7 Transcription Kit (Thermo Fisher Scientific) following the manufacturer’s protocol. The produce dsRNAs were resuspended in RNAase-free milliQ-water and stored at −20°C prior use.

### Fluorescence labeling of dsRNA

Fluorescence labeling of *SaMIF1*-dsRNA was performed using the HighYield T7 AF488 RNA Labeling Kit (Jena Bioscience, Germany) following the manufacturer’s instruction. Labeled *SaMIF1*-dsRNA_A488_ was used for the uptake experiments. For uptake analysis of small RNA, 21 nt *GAPDH*-dsRNA (provided in the kit) was labeled with Cy™ 3 utilizing the Silencer™ siRNA Labeling kit (ThermoFisher) according to the manufacturer’s instructions.

### Feeding of aphids on dsRNA supplemented artificial diet

The rearing method as described by Will et al. (2012) was used with minor modifications. The artificial diet (50 mM L-serine, 50 mM L-methionine, and 50 mM L-aspartic acid; pH 7.2) containing different sucrose concentrations was sealed between two layers of parafilm in a 2 cm diameter feeding tube, and one-day-old *Sa* nymphs were placed on the plates. The plates were covered with a feeding tube. The diet was prepared with RNase free water. For dsRNA feeding experiments, the dsRNA was mixed with the artificial diet. Ten synchronized nymphs with five replicates for each sample were used. Nymphs were placed at 22 □ under 65 % relative humidity, with a photoperiod of 16 h and a photon flux density of 125 μmol m^−2^ s^−1^.

### Application of dsRNA

Three-week old barley seedlings (each pot with three plants) were first sprayed with 0.02% Silwet-77, and 10 min later with 10 μg dsRNA solved in 500 μl deionized water. Controls were sprayed with 500 μL of deionized water. After spraying, the plants were infested with 50 *Sa* nymphs and stored in closed jars. Seven days later, the number of aphids was counted.

For treatment of Arabidopsis leaves, 19-day-old Arabidopsis SUC2::YFP seedlings grown in vertical plates were treated with 1 µL drop of nuclease free-water containing 20 µM dsRNA_Cy3_ at on the top of the leaf. Four leaves were treated. Confocal images were taken 5 days later.

### dsRNA delivery via the sucrose-aid method

Ten-day-old barley seedlings were transferred to the dark for 12 h. Leaves were detached and submerged with the basal end into 200 mM sucrose solution containing 20 µg / mL dsRNA for 24 h in the dark. Subsequently, the submerged parts of the leaves were cut and the top segment transferred to agar plates and used for aphid infestation. Thirty-two-day-old Arabidopsis leaves were cut and inserted with the petiole in nuclease-free water containing 20 µM dsRNA_Cy3_. For the sucrose-aid experiment, the solution was supplemented with 200 mM sucrose.

### Microscopy

Cross hand-cut sections of barley leaves were analyzed using a confocal laser-scanning microscopy (CLSM, Leica, TCS SP8, Germany). Green fluorescence of dsRNA_A488_ was detect by filter AF488 (λexc 494, λem 515 nm). The laser filter AF633 (λexc 631 nm, λem 642 nm) was used for the detection of red fluorescence, e.g. chloroplast, and autofluorescence of tissues). Arabidopsis leaves were visualized with the CLSM microscope (previously described) for fluorescence YFP (λ_exc_514 nm, λ_em_527 nm) and Cy3 (λ_exc_555nm, λ_em_569 nm). YFP was excited with the 514 nm laser (detection 519-551 nm) and Cy3 with the 561 nm laser (detection 566-635 nm).

## Discussion

We show here that members of the Macrophage migration inhibitor factor (MIF) protein family are necessary for the survival of the aphid *Sitobion avenae*. We found that *Sa* contains four *MIF* genes and that silencing of three of them, namely *SaMIF1, SaMIF2* and *SaMIF*3, leads to reduced aphid survival on artificial diet. This corroborates findings that MIFs, apart from its role in suppressing host immunity, also have an endogenous function in the aphid (Naessens et al. 2015). dsRNAs targeting individual *SaMIF* genes were effective at a concentration of 250 ng/µL. At lower concentration (125 ng/µL), only dsRNA directed against the *SaMIF1* transcript reduced target gene expression substantially, suggesting the possibility that *SaMIF1* could be a potential target candidate for aphid control by RNAi.

Functionally redundant *MIF* gene family members are wide-spread in eukaryotic genomes, which often hampers the analysis of gene families, due to functional redundancy (Jover-Gil et al. 2014; Martienssen and Irish 1999). For functional analysis, silencing of the entire set of paralogous genes at the same time is a straight forward approach. Simultaneous targeting of three out of the four known *SaMIF* genes using three *SaMIF* gene*-*specific dsRNAs caused a significant reduction in survival, when compared with the activity of a *GFP*-dsRNA that had no known target in *Sa* (Fig. 4). Interestingly, when applied in mixtures, *SaMIF*-dsRNAs had a synergistic effect as they affected survival of *Sa* in a concentration that showed no effects upon single delivery. Despite of this finding, overall our data suggest that *SaMIF1* is a candidate for aphid control and it is probably not required to take the other *SaMIF* genes into consideration.

In our experiments, different dsRNA delivery strategies were investigated to test the efficiency of RNAi-mediated control of insects. Oral feeding on artificial diet containing *SaMIF1-*dsRNA showed the highest mortality rate (Fig. 2) and concomitant downregulation of *SaMIF1* target transcripts (Fig. 3). In contrast, spraying *SaMIF1*-dsRNA onto leaves had no effect on the survival rate of nymphs fed on these leaves (Fig. S3). This result can be explained by the fact that the *SaMIF1*-dsRNA applied to the leaves did not reach the sieve-tubes in amounts sufficient to silence the *SaMIF1* target gene, though it cannot be excluded that spraying leaves with higher concentration of dsRNA would have an effect on aphid survival. Uptake of dsRNA via the leaf surface has been controversially discussed. Gogoi et al. (2017) published data showing that aphids take up, among others, a 588 bp long dsRNA from tomato leaves. It should be noted, however, that the dsRNA was applied by gently rubbing the solution onto the upper side of tomato leaflets that were previously carborundum-dusted. Subsequently, the treated leaves were thoroughly washed with 0.05% Triton X-100 for five times in 3 min intervals, showing that the dsRNA application method was rather harsh. In a report from the group of N. Mitter, dsRNA-mediated protection was obtained in tobacco against viral diseases, when leaves were spread with virus-specific dsRNA loaded on non-toxic, degradable, layered double hydroxide (LDH) clay nanosheets (Mitter et al. 2017). Once loaded on LDH, the dsRNA did not wash off, showed sustained release and could be detected on sprayed leaves even 30 days after application. Finally, it was recently reported that strong *GFP* transgene silencing was accomplished in tobacco and tomato by loading dsRNA into carbon dots (Schwartz et al. 2020). Chemical formulations not only enhance the uptake of RNA from leaves, but could also improve dsRNA penetration through the body wall of an insect, as shown for a polymer/detergent formulation that improves RNAi-induced mortality in the soybean aphid *Aphis glycines* (Zheng et al. 2019). In the light of these reports, more research should be focused on the dsRNA delivery strategies that might support more efficient use of RNAi-based plant protection.

Feeding of *Sa* on barley leaves immersed at the base in *SaMIF1-*dsRNA containing buffer also did not affect aphids’ survival nor could we detect an effect on *SaMIF1* target gene expression (Fig. S4). This setup was tested because we wanted to evaluate alternative experimental design for simple and rapid screening of potential dsRNA targets for aphid control. In agreement with a lack of effect on aphids, we could not detect fluorescence in phloem tissue when barley leaves had been submerged into fluorescence *SaMIF1*-dsRNA_A488_ solution. Instead, we detected fluorescence predominantly in the xylem parenchyma cells, mainly the contact cells (Fig. 5a-g). This is in agreement with earlier reports, where apical transport of exogenous dsRNA structurally is located within xylem structures (Dalakouras et al. 2018; Dalakouras et al. 2020). While the latter reports and our investigation support the view that dsRNA application onto leaves and via petioles results in the accumulation of RNA in the xylem, some reports challenge this generalized view: i. ODN-directed gene silencing in barley is mediated by passive vascular feeding of ODN through cut barley leaves using co-import of sucrose and negatively charged ODN molecules (Sun et al. 2005), resulting in ODN uptake into the leaf symplast and living cells. ii. back in 1990, the importance of the xylem-to-phloem pathway was underscored in a review that summarized work of the precedent two decades (van Bel 1990). Moreover, it is well accepted that exo/endocytosis processes are involved in the uptake of macromolecules from xylem tissue (Botha et al. 2008; Słupianek et al. 2019). iii. Turnip mosaic virus (TuMV) is a single-stranded RNA virus that can cause diseases in cruciferous plants. Viral RNA can move systemically through both phloem and xylem as membrane-associated complexes in plants (Wan et al. 2015).

Trafficking of vesicles carrying sRNAs has been observed between Arabidopsis and *Botrytis cinerea* (Cai et al. 2018). Exosomes derived from Tetraspanin-GFP Arabidopsis line could be visualized as fluorescent dots, demonstrating that sRNA transfer occurs via exosomes. Trafficking of sRNA in vesicular bodies might explain why fluorescence appears in a punctate manner in traversal leaf section (Fig. 5a-c). If the supplied RNA is being transferred from one cell to another via exo/endocytosis, the RNA would be packed into vesicles and thus fluorescence would be dotted.

The fact that we could not detect dsRNA_A488_ fluorescence in the barley phloem tissue led us to further experiments to substantiate a xylem-associated uptake of dsRNA. We repeated the RNA uptake experiments with the Arabidopsis *SUC2::4xYFP* reporter line, which is a more sensitive tool to distinguish between transport of solutes in xylem and phloem. When taken up by petioles, we detected 21 nt dsRNA_Cy3_ exclusively in the Arabidopsis xylem, and its signal did not overlap with the YFP fluorescence of the phloem (Fig. 6a-h). This result further substantiates the previous report showing that dsRNA uptake and its acropetal transport follows mainly the apoplastic route via the xylem (Dalakouras et al. 2018). It also shows that a possible exchange of dsRNA from xylem-to-phloem is not efficient enough to be detected in our fluorescence microscopy experiment nor to silence genes from aphids feeding on the phloem at least at the concentrations used here. Nevertheless and consistent with our finding, soaking roots in dsRNA solution conferred protection in rice and maize against stem-borer (Li et al. 2015), further showing the potential of the approach. We also used the Arabidopsis *SUC2::4xYFP* reporter line to follow the uptake of 21 nt dsRNA_Cy3_, upon dropping onto leaves (Fig. 6i-l). In agreement with the results from the barley spray experiments, we could detect fluorescence exclusively in the leaf xylem. While fluorescence imaging is sensitive and a well accpeted method, final proof of the absence of exogenously-applied dsRNA in the symplast, e.g in mesophyll cells and sieve-tubes in missing. In particular, the observation that virus-specific dsRNA, when scattered on leaves, is quite effective in reducing viral infections suggests that dsRNA - possibly assisted by physical means such as formulations and gentle leaf wounding - can lead to symplastic uptake of dsRNA.

## Declarations

### Ethics approval and consent to participate

Approved by all authors

## Authorship principles

All authors whose names appear on the submission

1. made substantial contributions to the conception or design of the work; or the acquisition, analysis, or interpretation of data; or the creation of new software used in the work;
2. drafted the work or revised it critically for important intellectual content;
3. approved the version to be published; and
4. agree to be accountable for all aspects of the work in ensuring that questions related to the accuracy or integrity of any part of the work are appropriately investigated and resolved.

## Authors’ contributions

S.L., J.I., M.J.L-C, A.v.B. and K-H.K. wrote the manuscript; S.L., J.I., and K-H.K. designed the study; M.M.P. prepared material for the experiments; S.L. and M.J.L-C conducted the experiments; K-H.K., J.I., M.J.L-C, S.L. and A.v.B. analyzed all data and drafted the figures. All authors commented and reviewed the final manuscript.

## Consent for publication

All authors declare consent of publication.

## Availability of data and material

All data generated or analyzed during this study are included in this published article [and its supplementary information files].

## Code availability (software application or custom code)

Not applicable.

## Acknowledgements

We thank Christina Birkenstock for technical assistance. The personnel of the HiLIFE Biocomplex unit at the University of Helsinki, a member of Instruct-ERIC Centre Finland, FINStruct, and Biocenter Finland, is also gratefully acknowledged. This work was funded in the cooperative German/French program by the Deutsche Forschungsgemeinschaft (DFG) and the Agence Nationale de la Recherche (ANR) to KHK (Ko 1208/25-1) as well as in the DFG program FOR5116 to KHK and Academy of Finland (grant #331627) to MMP. SL was supported by the China Scholarship Council.

## Supplement Figures

**Fig. S1.**
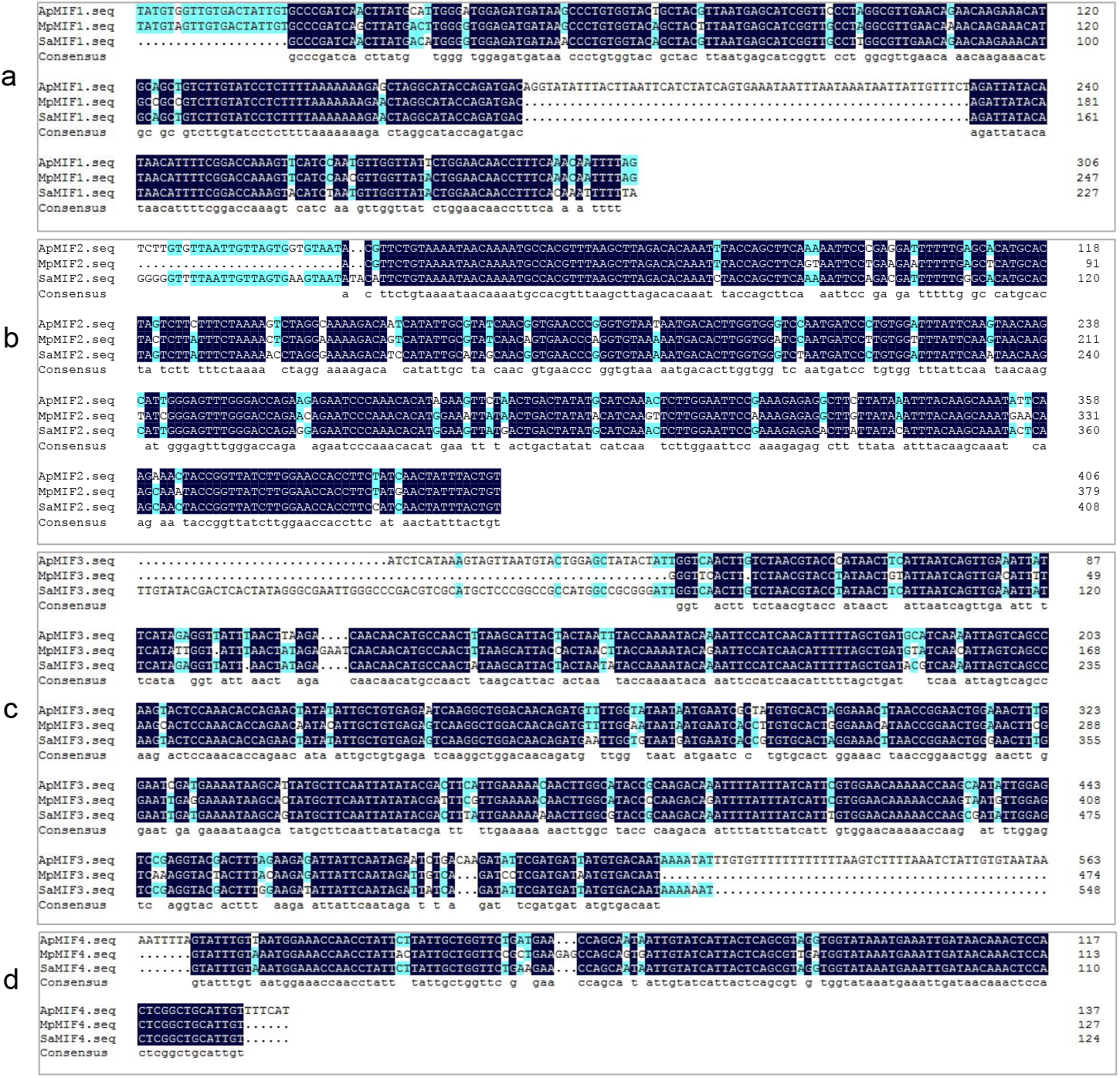
Prediction of partial *Sitobion avenae SaMIF* sequences using available sequence data of *Acyrthosiphon pisum* (*Ap*) and *Myzus persicae* (*Mp*). a, *SaMIF1*; b, *SaMIF2* c, *SaMIF3*; d, *SaMIF4*. Dark blue boxes denote homology. Sequences are deduced from *MpMIF1* (GenBank: KP218519), *MpMIF3* (GenBank: KR136352), *MpMIF4* (GenBank: KR136353), *ApMIF1* (LOC100161225), *ApMIF3* (LOC100144890) and *ApMIF4* (LOC100162394). The sequence of *SaMIF2* (JK723326) was published in Dubreuil et al. (2014).

**Fig. S2.**
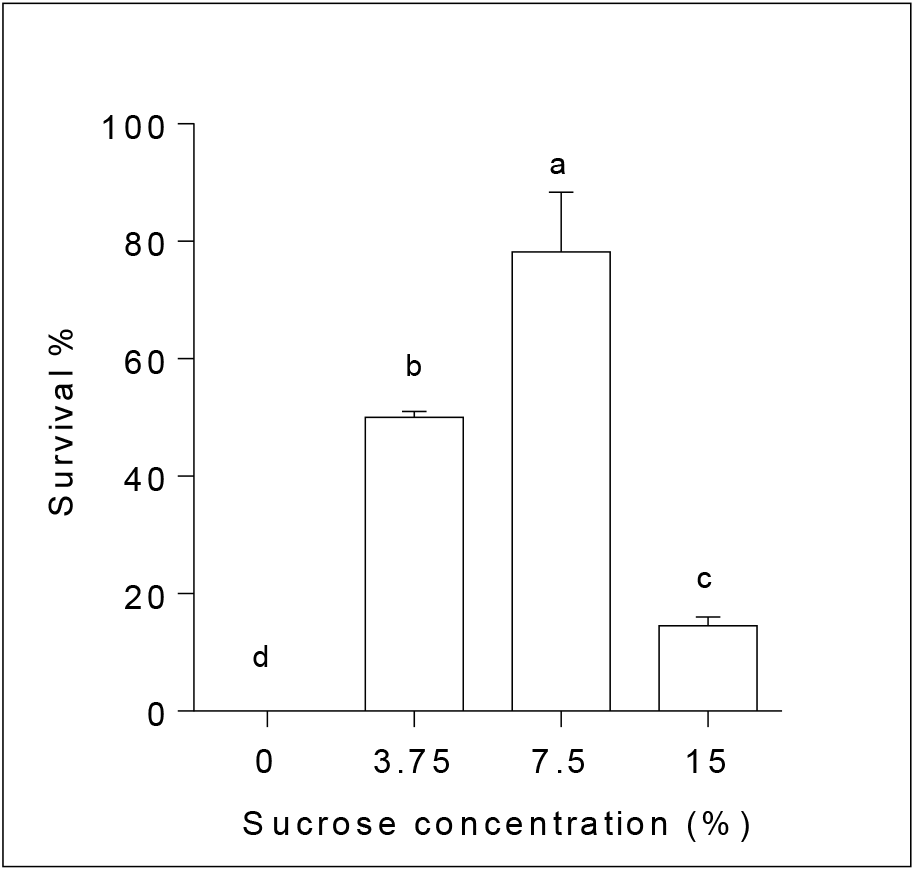
Survival of *Sitobion avenae* nymphs on artificial diet after four days of feeding. Artificial diet was supplemented with different sucrose concentrations. The most suitable concentration is 7.5% (w/v), which corresponds to 218 mM sucrose. Bars represent means (± SD) of three biological replicates. Survival data were evaluated by Kaplan-Meier analysis and log-rank test. Different letters indicate significant differences at p < 0.001.

**Fig. S3.**
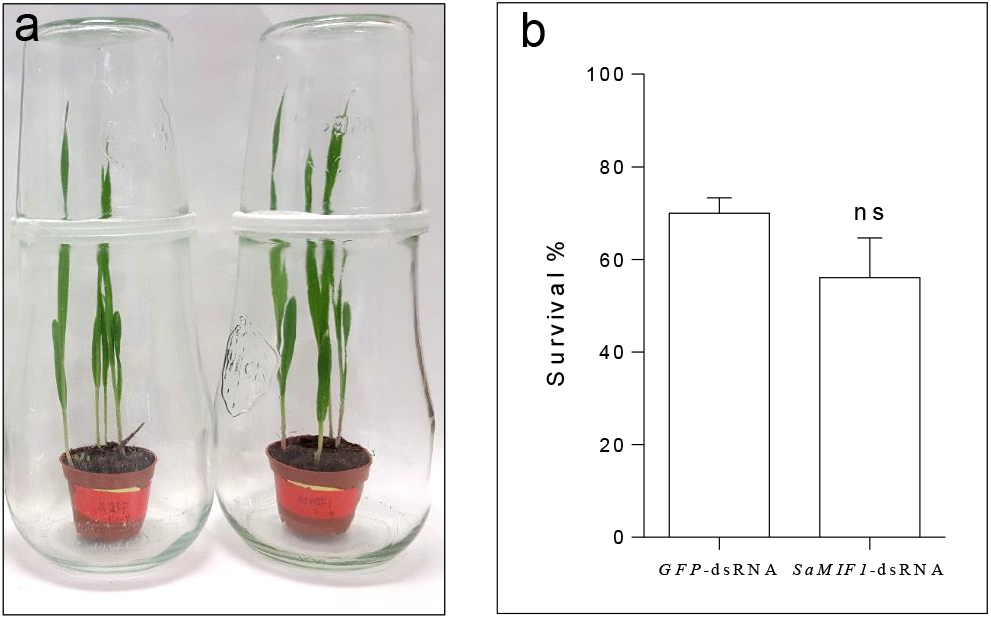
Survival analysis of aphids fed on plants sprayed with dsRNA after 7 days. **a**, experimental design: fifty synchronous one-day-old nymphs were kept on barley plants sprayed with *GFP-*dsRNA (20 ng/µL) or *SaMIF1-*dsRNA (20 ng/µL). The infested plants were kept in glass jars in a climate chamber with a 16 h photoperiod (260 µmol*/m*^*2*^*·s*^*-1*^) at 22°C/18°C (light/dark) with 65% relative humidity. **b**, aphids survival was monitored on day seven after the onset of feeding on sprayed plants. Bars represent mean values ± SD of three independent experiments. “ns” indicates no significant differences (p > 0.05).

**Fig. S4.**
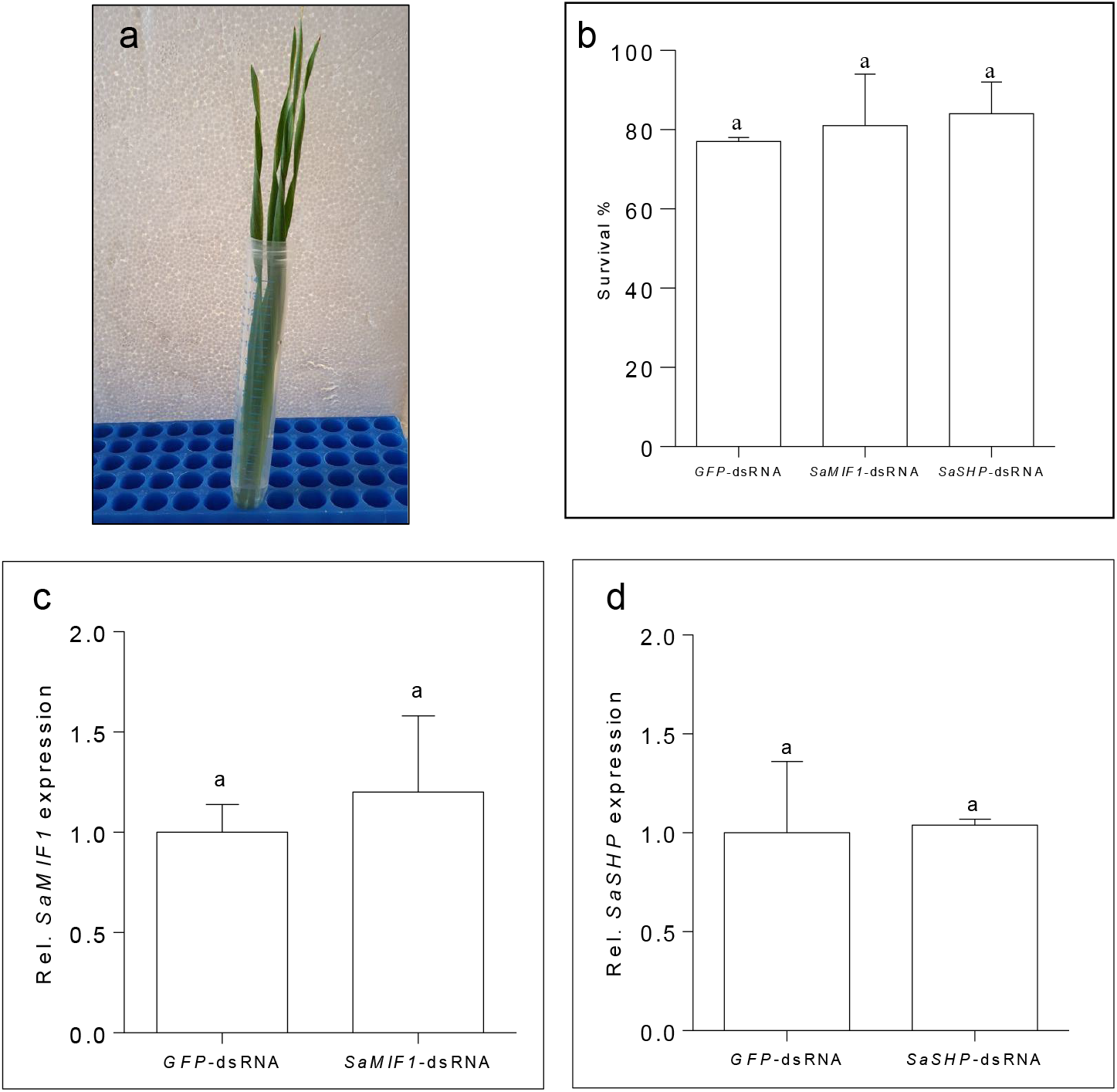
Survival and target gene expression of *Sitobion avenae* nymphs on detached barley leaves, the cut base immerged in *SaMIF1*-dsRNA or *SaSHP*-dsRNA, respectively. **a**, design of sucrose-aided dsRNA delivery method, cut bases of barley leaves were immerged in 200 mM sucrose containing 20 µg dsRNA. **b**, aphid survival monitored after seven days of feeding on barley leaves supplied with dsRNA solution. Bars represent mean values ± SD of three independent experiments. Relative expression level of *SaMIF1* (**c**) and *SaSHP* (**d**) transcripts normalized to *Rpl*27 analyzed by RT-qPCR. Bars represent means (± SD) from two independent replicates. “ns” indicates no significant differences at p > 0.05.

**Fig. S5.**
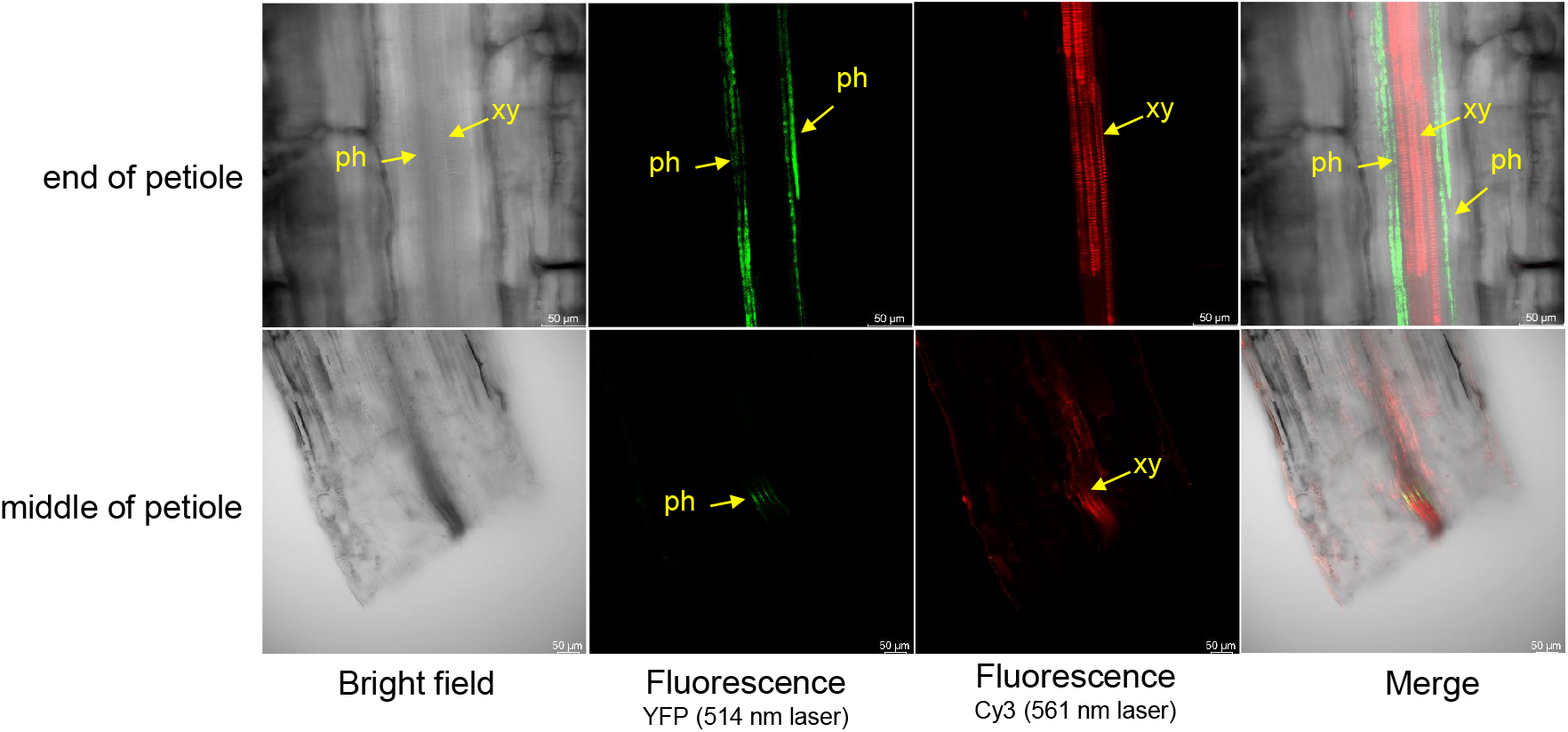
Feeding of 21 nt dsRNA_Cy3_ via *Arabidopsis thaliana* petioles. Confocal images of petiole segments of the reporter line *SUC2::YFP*. Cut petioles were immerged in 9.6 µL of a 200 mM sucrose solution containing 20 µM dsRNA_Cy3_ for 24 h. Images were taken with a confocal microscope from different segments of the petiole. The red color, which is restricted to the xylem vessels represents Cy3 (λ_exc_555nm, λ_em_569 nm) and the green color represents the phloem-based YFP fluorescence (λ_exc_514 nm, λ_em_527 nm). xy, xylem; ph, phloem.

