## supplemental tables for "Evaluation of dsRNA delivery methods for targeting macrophage migration inhibitory factor MIF in RNAi-based aphid control"

**Supplementary**

**Material and Methods**

Table S1. Primers used in the study

| Primer name | Primer sequence | Application |
|---|---|---|
| SaMIF1_F | AATATGCCTCATTTCCGTTTGGA | degenerate primer |
| SaMIF1_R | CCTAAAATTGTTTGAAAGGTTGTT | degenerate primer |
| *Sa*MIF1 _T7_F | TAATACGACTCACTATAGGGAGAAATATGCCT CATTTCCGTTTGG | dsRNA synthesis[1] |
| *Sa*MIF1 _T7_R | TAATACGACTCACTATAGGGAGATGTTCTGTT CAACGCCAAGG | dsRNA synthesis[1] |
| qSaMIF1_F | CCTTGGCGTTGAACAGAACA | qPCR |
| qSaMIF1_R | GATGTACTTTGGTCCGAAAATGT | qPCR |
| S*a*MIF2_RNAi_F | TAATACGACTCACTATAGGGAGATGCCACGTT TAAGCTTAGACACAA | dsRNA synthesis[1] |
| SaMIF2_RNAi_R | TAATACGACTCACTATAGGGAGAACCGGTAGT TGCTTGAGTATTTGC | dsRNA synthesis[1] |
| SaMIF2_T7 | TAATACGACTCACTATAGGGTGCCACGTTTAA GCTTAGACAC | dsRNA synthesis[2] |
| SaMIF2_phi6 | GGAAAAAAAACCGGTAGTTGCTTGAGTATTT G | dsRNA synthesis[2] |
| qSaMIF2_F | GCCACGTTTAAGCTTAGACA | qPCR |
| qSaMIF2_R | GATTCTCCTCTGGTCCCAAA | qPCR |
| SaMIF3-2_F | GGTCAACTTGTCTAACGTACC | degenerate primer |
| SaMIF3-2_R | TTTTATTGTCACATAATCATCGA | degenerate primer |
| SaMIF3_T7_F | TAATACGACTCACTATAGGGAGAGCCAAGTAC TCCAAACACCA | dsRNA synthesis[1] |
| SaMIF3_T7_R | TAATACGACTCACTATAGGGAGATGATAACT AAAATTTGTCTTGCGGT | dsRNA synthesis[1] |
| qSaMIF3_F | ACAACAACATGCCAACTATAAGC | qPCR |
| qSaMIF3_R | TCCAGCCTTGACTCTCACAG | qPCR |
| SaMIF4_F | GTATTTGTAAATGGAAACCAACCTA | degenerate primer |
| SaMIF4_R | ACAATGCAGCCGAGTGGA | degenerate primer |
| dsEGFP_T7_F | CCCTTTAATACGACTCACTATAGGGAGAACCA CATGAAGCAGCACGAC | dsRNA synthesis[1] |
| dsEGFP_T7_R | CCCTTTAATACGACTCACTATAGGGAGAGTCC ATGCCGAGAGTGATC | dsRNA synthesis[1] |
| EGFP_T7 | TAATACGACTCACTATAGGGACCACATGAAGC AGCACG | dsRNA synthesis[2] |
| EGFP_phi6 | GGAAAAAAAGTCCATGCCGAGAGTGATC | dsRNA synthesis[2] |
| *Sa*SHP _T7_F | TAATACGACTCACTATAGGGAGAAGTCGCTG TAATCGGTGCTGATG | dsRNA synthesis |
| *Sa*SHP _T7_R | TAATACGACTCACTATAGGGAGAACGCGTCG ATGTCAAACAGGC | dsRNA synthesis |
| SaSHP_T7 | TAATACGACTCACTATAGGGAGTCGCTGTAAT CGGTGCTG | dsRNA synthesis[2] |
| SaSHP_phi6 | GGAAAAAAAACGCGTCGATGTCAAACAG | dsRNA synthesis[2] |
| qSaSHP_F | ACACCAACATACATTGACCAGC | qPCR |
| qSaSHP_R | TGGGTTTCTGTACATTGGGAAT | qPCR |
| qRpl27_F | CCGAAAAGCTGTCATAATGAAGACC | qPCR |
| qRpl27_R | GGTGAAACCTTGTCTACTGTTACATCTTG | qPCR |

[1] Primers used for the amplification of target sequence for subsequent dsRNA production using MEGAscript T7 Transcription Kit (Thermo Fisher Scientific).

[2] Pimers used for the amplification of the target sequence for subsequent dsRNA production using T7 and phi6 RNA polymerases.

Table S2. Target gene sequences

| **Sequences used for RNA silencing** |
|---|
| ***Sa*MIF1 (223 bp)**<br>AATATGCCTCATTTCCGTTTGGAAACTAATGTTTCAAAGTCTAAAGTAACACCAGAATTATTGAAA<br>AAGCTTTCTGCAGCTGTGGCCAAAACTCTTGGAAAACCGGAATCGTATGTAGTTGTGACTGTTGT<br>GCCCGATCAACTTATGACATGGGGTGGAGATGATAAACCCTGTGGTACAGCTACGTTAATGAGCAT<br>CGGTTGCCTTGGCGTTGAACAGAACA |
| **SaMIF2 (323 bp)**<br>**TGCCACGTTTAAGCTTAGACAC**AAATCTACCAGCTTCAAAAATTCCAGACGATTTTTTGGGCAC<br>ATGCACTAGTCTTATTTCTAAAAACCTAGGGAAAAGACATCCATATTGCATAGCAACGGTGAAC<br>CCGGGTGTAAAAATGACACTTGGTGGGTCTAATGATCCCTGTGGATTTATTCAAATAACAAGCAT<br>TGGGAGTTTGGGACCAGAGGAGAATCCCAAACACATGGAAGTTATGACTGACTATATGCATCAA<br>ACTCTTGGAATTCCGAAAGAGAGACTTATTATACATTTACAAGCAAATACTCAAGCAACTACCG<br>GT |
| **SaMIF3 (212 bp)**<br>GCCAAGTACTCCAAACACCAGAACTATATATTGCTGTGAGAGTCAAGGCTGGACAACAGATGAAT<br>TGGTATAATGATGAATCACCGTGTGCACTAGGAAACTTAACCGGAACTGGAAACTTTGGAATTGAT<br>GAAAATAAGCAGTATGCTTCAATTATATACGACTTTATTGAAAAAAAACTTGGCGTACCGCAAGAC<br>AAATTTTAGTTATCA |
| **SaSHP (470 bp)**<br>AGTCGCTGTAATCGGTGCTGATGACTGTTGGAAAGAGTATATGGTGTTAGCTGTTATCAACAAGT<br>ATGACAACTTCTTTGGTGGAAATGACCATATTATTTGGGTAGTGACGAGAGATGTTAACCGTAAT<br>TGGTCTACTTATGATAAAGCATACAACGATATTAAGGAAAGTGGTCTCTGCCCTAACATTTTGGT<br>AAGTGTTGATCATTCGTTTGAACCCATGACAGGACCTTCAATAGCAGTACCTTCAATGGAACCAA<br>GTGTGGCAGTACCTTCAATGGCACCAACTATGCCTGGTGATATCGATGGCATGGTACAAACAAC<br>GTCTGTCTCTACAACATCAGCAACGAAATCAATAAGTACCGACGGCGATACTGCTGTAACTTCAT<br>CATCCACTTCAACAACTACGACATCGACTGTGATAATTGATAAAAGTGATGATTTCTCTTGCCTG<br>TTTGACATCGACGCGT |
| **GFP (476 bp)**<br>ACCACATGAAGCAGCACGACTTCTTCAAGTCCGCCATGCCCGAAGGCTACGTCCAGGAGCGCAC<br>CATCTTCTTCAAGGACGACGGCAACTACAAGACCCGCGCCGAGGTGAAGTTCGAGGGCGACACC<br>CTGGTGAACCGCATCGAGCTGAAGGGCATCGACTTCAAGGAGGACGGCAACATCCTGGGGCACA<br>AGCTGGAGTACAACTACAACAGCCACAACGTCTATATCATGGCCGACAAGCAGAAGAACGGCATC<br>AAGGTGAACTTCAAGATCCGCCACAACATCGAGGACGGCAGCGTGCAGCTCGCCGACCACTACC<br>AGCAGAACACCCCCATCGGCGACGGCCCCGTGCTGCTGCCCGACAACCACTACCTGAGCACCCA<br>GTCCGCCCTGAGCAAAGACCCCAACGAGAAGCGCGATCACATGGTCCTGCTGGAGTTCGTGACC<br>GCCGCCGGGATCACTCTCGGCATGGAC |
